# *Arid1b* haploinsufficiency in pyramidal neurons causes cellular and circuit changes in neocortex but is not sufficient to produce behavioral or seizure phenotypes

**DOI:** 10.1101/2024.06.04.597344

**Authors:** Alec H. Marshall, Meretta A. Hanson, Danielle J. Boyle, Devipriyanka Nagarajan, Noor Bibi, Julie Fitzgerald, Emilee Gaitten, Olga N. Kokiko-Cochran, Bin Gu, Jason C. Wester

**Author notes:** These authors contributed equally (Alec H. Marshall, Meretta A. Hanson, Danielle J. Boyle).

## Abstract

*Arid1b* is a high confidence risk gene for autism spectrum disorder that encodes a subunit of a chromatin remodeling complex expressed in neuronal progenitors. Haploinsufficiency causes a broad range of social, behavioral, and intellectual disability phenotypes, including Coffin-Siris syndrome. Recent work using transgenic mouse models suggests pathology is due to deficits in proliferation, survival, and synaptic development of cortical neurons. However, there is conflicting evidence regarding the relative roles of excitatory projection neurons and inhibitory interneurons in generating abnormal cognitive and behavioral phenotypes. Here, we conditionally knocked out either one or both copies of *Arid1b* from excitatory projection neuron progenitors and systematically investigated the effects on intrinsic membrane properties, synaptic physiology, social behavior, and seizure susceptibility. We found that disrupting *Arid1b* expression in excitatory neurons alters their membrane properties, including hyperpolarizing action potential threshold; however, these changes depend on neuronal subtype. Using paired whole-cell recordings, we found increased synaptic connectivity rate between projection neurons. Furthermore, we found reduced strength of excitatory synapses to parvalbumin (PV)-expression inhibitory interneurons. These data suggest an increase in the ratio of excitation to inhibition. However, the strength of inhibitory synapses from PV interneurons to excitatory neurons was enhanced, which may rebalance this ratio. Indeed, *Arid1b* haploinsufficiency in projection neurons was insufficient to cause social deficits and seizure phenotypes observed in a preclinical germline haploinsufficient mouse model. Our data suggest that while excitatory projection neurons likely contribute to autistic phenotypes, pathology in these cells is not the primary cause.

## INTRODUCTION

Autism spectrum disorder (ASD) presents with diverse symptoms that include deficits in social behavior, communication, sensory processing, and learning and memory. It remains unresolved how or why these converge in ASD patients, but a prominent hypothesis is that they occur due to an imbalance between excitation and inhibition within cortical circuits (Sohal and Rubenstein, 2019). To find potential sources of pathological imbalance, there has been great effort to characterize changes in neuronal excitability and synaptic physiology in a broad range of ASD mouse models (Contractor et al., 2021; Ford et al., 2022; Monday et al., 2023). However, determining the pathophysiological mechanisms that cause ASD has remained elusive. This is due in part to the great diversity of neuronal subtypes in the cortex and the plasticity of the synapses between them.

Monogenetic mouse models of ASD are a powerful tool to investigate cell-type-specific mechanisms. Recently, several labs have focused on a model of haploinsufficiency for *Arid1b* (Moffat et al., 2021a). *Arid1b* is a high confidence risk gene for ASD (Hoyer et al., 2012; De Rubeis et al., 2014; Satterstrom et al., 2020), and mutations are responsible for Coffin-Siris syndrome (Santen et al., 2012; van der Sluijs et al., 2019). ARID1B is a subunit of a chromatin remodeling complex expressed in progenitors of both excitatory and inhibitory neurons (Son and Crabtree, 2014; Jung et al., 2017; Moffat et al., 2021b; Moffat et al., 2021a). In mice, *Arid1b* haploinsufficiency leads to several phenotypes consistent with ASD, including social and cognitive deficits and repetitive behaviors (Celen et al., 2017; Jung et al., 2017; Shibutani et al., 2017; Ellegood et al., 2021; Kim et al., 2022). ARID1B’s function is broadly associated with neuronal proliferation and differentiation (Moffat et al., 2021b; Pagliaroli and Trizzino, 2021), but it remains unclear how mutations cause ASD.

Studies in mice provide evidence that *Arid1b* haploinsufficiency in excitatory pyramidal neurons contributes to phenotypic abnormalities (Ka et al., 2016; Jung et al., 2017; Moffat et al., 2021b; Kim et al., 2022). However, as is common in ASD research, the data are conflicting and the role of pyramidal neurons relative to inhibitory interneurons remains unclear. Early work using *in utero* delivery of short hairpin RNA to knockdown *Arid1b* in layer 2/3 pyramidal neurons found reductions in dendritic complexity and spine density (Ka et al., 2016). Subsequently, multiple groups generated transgenic mice with either whole-body *Arid1b* haploinsufficiency as a preclinical model, or a floxed *Arid1b* allele for conditional knockout from either excitatory or inhibitory neuronal subtypes. In a preclinical model, Kim et al. (2022) found reduced frequency of miniature excitatory postsynaptic currents (mEPSCs) in layer 2/3 pyramidal neurons, consistent with the earlier observation of disrupted excitatory synapse formation (Ka et al., 2016). In contrast, Jung et al. (2017) did not find reduced mEPSC frequency in pyramidal neurons; instead, they provide strong evidence for pathology in inhibitory interneurons and reduced cortical inhibition. Finally, Moffat et al. (2021b) conditionally knocked out both copies of *Arid1b* from pyramidal neuron progenitors. They found reduced pyramidal neuron proliferation, but complete loss of *Arid1b* from pyramidal neurons could only replicate a subset of ASD phenotypes observed in the preclinical *Arid1b* haploinsufficient model. Thus, it remains unclear if cell-autonomous pathology in pyramidal neurons is a major driver of ASD phenotypes in *Arid1b* haploinsufficiency.

Here, we used transgenic mice to manipulate *Arid1b* expression conditionally in pyramidal neurons. We found that *Arid1b* disfunction affects pyramidal neuron differentiation and intrinsic membrane properties dependent on subtype. Thus, pathology in pyramidal neurons is not uniform across cells. Furthermore, we found changes in excitatory synaptic connectivity and strength that suggest imbalanced excitation-to-inhibition. However, these were countered by potentiation of inhibitory synapses onto pyramidal neurons. Ultimately, *Arid1b* haploinsufficiency in pyramidal neurons is insufficient to produce behavioral and seizure phenotypes observed in the preclinical mouse model. Our data are consistent with the hypothesis that pathology in pyramidal neurons contributes to, but does not drive, ASD phenotypes in *Arid1b* haploinsufficiency (Jung et al., 2017; Moffat et al., 2021b).

## RESULTS

To determine if ARID1B disfunction in pyramidal neurons (PNs) causes pathology in cellular and synaptic physiology, we crossed Emx1-IRES-Cre mice (Gorski et al., 2002) to a transgenic line in which loxP sites flank exon 5 of the *Arid1b* allele (Celen et al., 2017). This allowed us to conditionally knockout one or both copies of *Arid1b* from PNs during embryogenesis. Throughout, we term these mice PN *Arid1b*(+/-) and PN *Arid1b*(-/-), respectively. We performed our physiology experiments in slices of primary visual cortex (V1), where neuronal cell-types and circuits are well studied. To reveal potential pathological changes in cellular and synaptic physiology between conditions, our strategy was to compare neurons at a stable development stage. Thus, we used 5 – 8-week-old mice to study mature neurons after the critical period for plasticity (Gordon and Stryker, 1996; Levelt and Hubener, 2012). Our focus was on the effect of *Arid1b* haploinsufficiency because of its relevance as a preclinical model. However, for a subset of initial experiments, we used PN *Arid1b*(-/-) mice to supplement previous data reported by Moffat et al. (2021b).

### Pyramidal neurons in layer 2/3 can be separated into two groups by firing type that are differentially vulnerable to *Arid1b* disfunction

PNs in layer 2/3 express unique molecular markers, suggesting diversity among this cell population (Molyneaux et al., 2009). Indeed, recent work found that layer 2/3 PNs in V1 generate differential firing patterns observed in vivo that can be linked to their unique electrophysiological properties recorded in brain slices (Wei et al., 2023). Thus, we first recorded a large population of PNs in control mice (n = 81 cells) to determine if they could be separated into different types that might be uniquely impacted by loss of *Arid1b*.

We found that layer 2/3 PNs could be distinguished by their spike frequency adaptation during constant amplitude current steps (**Fig. 1A**). We identified two broad PN subtypes that we refer to as continuous adapting (CA) (**Fig. 1A, left**) and fast adapting (FA) (**Fig. 1A, right**). PNs with FA responses in layer 2/3 of V1 were recently identified by two groups (Gouwens et al., 2019; Wei et al., 2023) and can be found in the Allen Cell Types Database (http://celltypes.brain-map.org/). However, the physiology of CA and FA subtypes in layer 2/3 have not been previously compared in detail. As expected, CA cells had a higher firing rate than FA cells in response to constant amplitude current steps at twice the current threshold for action potential initiation (**Fig. 1B**). CA cells also demonstrated a greater afterhyperpolarization amplitude following the first spike (**Fig. 1C**). However, the voltage threshold for spike initiation (**Fig. 1D**) and the spike half-width at half-height (**Fig. 1E**) were not different between CA and FA cells. CA and FA cells also had different subthreshold membrane properties: CA cells had a higher input resistance (**Fig. 1F**) and a larger estimated membrane capacitance (**Fig. 1G**). However, CA and FA cells had equivalent voltage sag in response to hyperpolarizing current pulses (**Fig. 1H**), suggesting similar expression of HCN channels.

**Figure 1.**
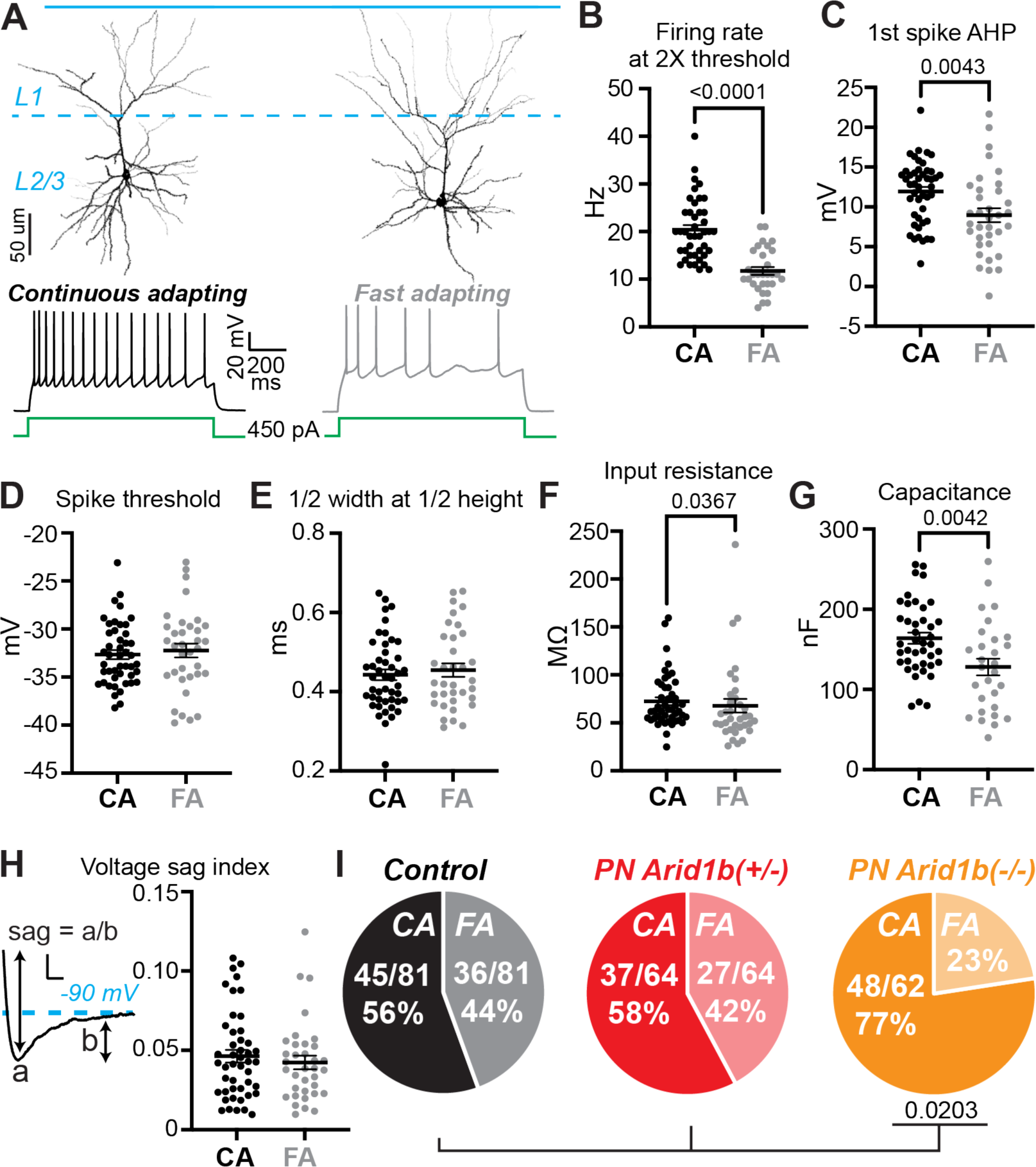
Two types of pyramidal neurons in layer 2/3 of V1 can be identified by intrinsic membrane properties and are differentially affected by loss of *Arid1b*. **A)** Example of continuous adapting (CA) and fast adapting (FA) PN types in V1 layer 2/3. Reconstructions of dendrites traced from biocytin-filled cells. **B)** Firing rate at twice action potential threshold. CA n = 42, FA n = 30; Mann-Whitney U test. **C)** Afterhyperpolarization amplitude after the first action potential. CA n = 46, FA n = 35; unpaired t test. **D)** Action potential voltage threshold. CA n = 46, FA n = 35; p = 0.6156, t test. **E)** Half-width at half-height for the first action potential. CA n = 46, FA n = 35; p = 0.7364, Mann-Whitney test. **F)** Input resistance. CA n = 46, FA n = 35; Mann-Whitney test. **G)** Membrane capacitance. CA n = 41, FA n = 29; p = 0.6156, unpaired t test. **H)** Hyperpolarization-induced membrane potential voltage sag. Inset demonstrates the calculation of voltage sag index. CA n = 46, FA n = 35; p = 0.5855, Mann-Whitney U test. Scale bar: 2 mV, 20 ms. **I)** Proportions of CA and FA PN types across mouse conditions. Chi-squared test (df = 2).

In control mice, we found that just over half of recorded PNs were of the CA subtype (45/81, 56%), while the remainder were classified as FA (36/81, 44%) (**Fig. 1I, left**). In PN *Arid1b*(+/-) mice these proportions were unchanged (**Fig. 1I, middle**), suggesting *Arid1b* haploinsufficiency does not differentially alter the proliferation of these subtypes. However, complete loss of *Arid1b* resulted in a striking reduction in the number of FA cells relative to CA cells (**Fig. 1I, right**). Recently, Moffat et al. (2021b) used a similar transgenic strategy to knockout both copies of *Arid1b* from PNs and found that loss of Arid1b resulted in reduced density of CUX1+ cells in layer 2/3. Our data are consistent with their findings but suggest that a specific subpopulation of these PNs, identified by FA firing type, is uniquely vulnerable.

Next, we investigated if *Arid1b* disfunction differentially affects the electrophysiological properties of CA and FA PN cell-types. We began with CA PNs recorded from control, PN *Arid1b*(+/-), and PN *Arid1b*(-/-) mice because we were able to observe a sufficient number of these cells to make comparisons among each condition. Strikingly, we found that haploinsufficiency for *Arid1b* had no effect on CA PN membrane properties and that complete loss resulted in few changes. In PN *Arid1b*(-/-) mice, the first spike width was decreased (**Fig. 2A, B**) and the first spike afterhyperpolarization increased (**Fig. 2A, C**). However, the voltage threshold for spike initiation (**Fig. 2D**), the firing rate at twice spike threshold (**Fig. 2E**), input resistance (**Fig. 2F**), membrane capacitance (**Fig. 2G**), and voltage sag (**Fig. 2H**) were not different among conditions. Next, we compared FA PNs recorded in control and PN *Arid1b*(+/-) mice. In contrast to CA PNs, *Arid1b* haploinsufficiency in FA PNs was sufficient to cause a reduction in spike width (**Fig. 2I, J**). Furthermore, FA cells in PN *Arid1b*(+/-) mice had a hyperpolarized voltage threshold for spike initiation (**Fig. 2I, K**). However, the first spike afterhyperpolarization (**Fig. 2L**) and firing rate at twice spike threshold (**Fig. 2M**) were not affected. Finally, although input resistance was not affected (**Fig. 2N**), FA cells in PN *Arid1b*(+/-) mice had increased membrane capacitance (**Fig. 2O**) and increased voltage sag (**Fig. 2P**).

**Figure 2.**
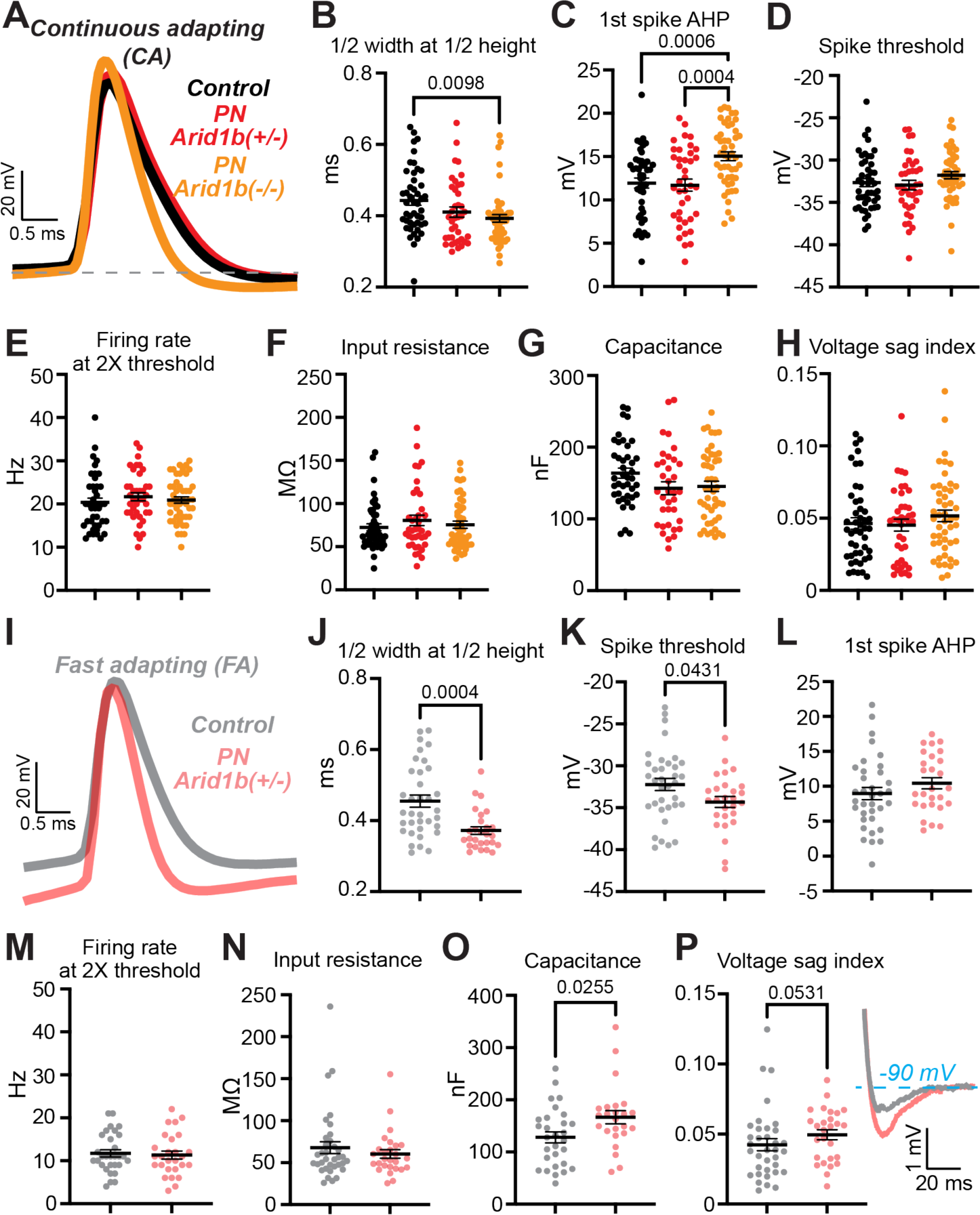
*Arid1b* haploinsufficiency preferentially disrupts the membrane properties of fast adapting pyramidal neurons in layer 2/3. **A)** Examples of first evoked action potential across conditions for the CA PN type. Note narrower spike with larger afterhyperpolarization in PN *Arid1b*(-/-) mice, but normal spike in PN *Arid1b*(+/-) mice. **B)** Half-width at half-height for the first action potential. CA Control n = 46, PN *Arid1b*(+/-) n = 39, PN *Arid1b*(-/-) n = 48; Kruskal-Wallis test: H(2) = 9.239, p = 0.0099; Dunn’s post hoc: Control vs. PN *Arid1b*(+/-) p = 0.1097; PN *Arid1b*(+/-) vs. PN *Arid1b*(-/-) p > 0.9999. **C)** Afterhyperpolarization amplitude after the first action potential. CA Control n = 46, PN *Arid1b*(+/-) n = 39, PN Arid1b(-/-) n = 48; one-way ANOVA: F(2,130) = 10.2, p < 0.0001; Tukey’s post hoc: Control vs. PN *Arid1b*(+/-) p = 0.9581. **D)** Action potential voltage threshold. CA Control n = 46, PN *Arid1b*(+/-) n = 39, PN *Arid1b*(-/-) n = 48; one-way ANOVA: F(2,130) = 1.628, p = 0.2003. **E)** Firing rate at twice action potential threshold. CA Control n = 42, PN *Arid1b*(+/-) n = 39, PN *Arid1b*(-/-) n = 47; Kruskal-Wallis test: H(2) = 1.806, p = 0.4053. **F)** Input resistance. CA Control n = 46, PN *Arid1b*(+/-) n = 39, PN *Arid1b*(-/-) n = 48; Kruskal-Wallis test: H(2) = 0.6066, p = 0.7384. **G)** Membrane capacitance. CA Control n = 41, PN *Arid1b*(+/-) n = 35, PN *Arid1b*(-/-) n = 47; Kruskal-Wallis test: H(2) = 4.528, p = 0.1039. **H)** Hyperpolarization-induced membrane potential voltage sag. CA Control n = 46, PN *Arid1b*(+/-) n = 39, PN *Arid1b*(-/-) n = 48; Kruskal-Wallis test: H(2) = 1.519, p = 0.4678. **I)** Examples of first evoked action potential between control and PN *Arid1b*(+/-) conditions for the FA PN type. Note hyperpolarized action potential voltage threshold and narrow spike width. **J)** Half-width at half-height for the first action potential. FA Control n = 35, PN *Arid1b*(+/-) n = 27; Mann-Whitney test. **K)** Action potential voltage threshold. FA Control n = 35, PN *Arid1b*(+/-) n = 27; t test. **L)** Afterhyperpolarization amplitude after the first action potential. FA Control n = 35, PN *Arid1b*(+/-) n = 27; p = 0.2346, t test. **M)** Firing rate at twice action potential threshold. FA Control n = 30, PN *Arid1b*(+/-) n = 27; p = 0.7510, t test. **N)** Input resistance. FA Control n = 35, PN *Arid1b*(+/-) n = 27; p = 0.7780, Mann-Whitney test. **O)** Membrane capacitance. FA Control n = 29, PN *Arid1b*(+/-) n = 25; Mann-Whitney test. **P)** Hyperpolarization-induced membrane potential voltage sag. FA Control n = 35, PN *Arid1b*(+/-) n = 27; Mann-Whitney test.

Collectively, these data suggest that in layer 2/3, FA PNs are selectively vulnerable to *Arid1b* disfunction. Complete loss of *Arid1b* reduces their density in mature mice and haploinsufficiency alters their electrophysiological properties, including spike threshold. Although the changes are subtle, collectively they may be sufficient to disrupt information flow through the cortex.

### The intrinsic membrane properties of pyramidal tract-type pyramidal neurons in layer 5 are also affected by *Arid1b* haploinsufficiency

Although layer 2/3 PNs include different subtypes, they are all broadly part of the same intratelencephalic projection class (Harris and Shepherd, 2015). Thus, to determine if the effects of *Arid1b* haploinsufficiency on PN electrophysiological properties are dependent on PN class, we next recorded from pyramidal tract-type (PT) PNs in layer 5. To target these cells in V1, we injected red retrograde tracer into the superior colliculus (**Fig. 3A, top**). Labeled PT-type PNs had a thick-tufted morphology (**Fig. 3A, bottom**) and commonly demonstrated initial burst firing (**Fig. 3B**), as described previously (Morishima and Kawaguchi, 2006; Harris and Shepherd, 2015). Interestingly, *Arid1b* haploinsufficiency altered a similar set of membrane properties in layer 5 PT-type PNs as observed in layer 2/3 FA PNs. Like layer 2/3 PNs, *Arid1b* haploinsufficient PT-type PNs had a hyperpolarized voltage threshold for spike initiation (**Fig. 3B, bottom and 3C**) and narrower spike width (**Fig. 3D**), but the first spike afterhyperpolarization (**Fig. 3E**) and firing rate at twice spike threshold (**Fig. 3F**) were not affected. Furthermore, we observed an increase in voltage sag in PT-type PNs (**Fig. 3G**). However, we found no change in input resistance (**Fig. 3H**) or membrane capacitance (**Fig. 3I**). Thus, *Arid1b* haploinsufficiency causes a common set of membrane property changes across PN classes and cortical layers. This may broadly disrupt cortical information processing in diverse circuits dedicated to unique tasks.

**Figure 3.**
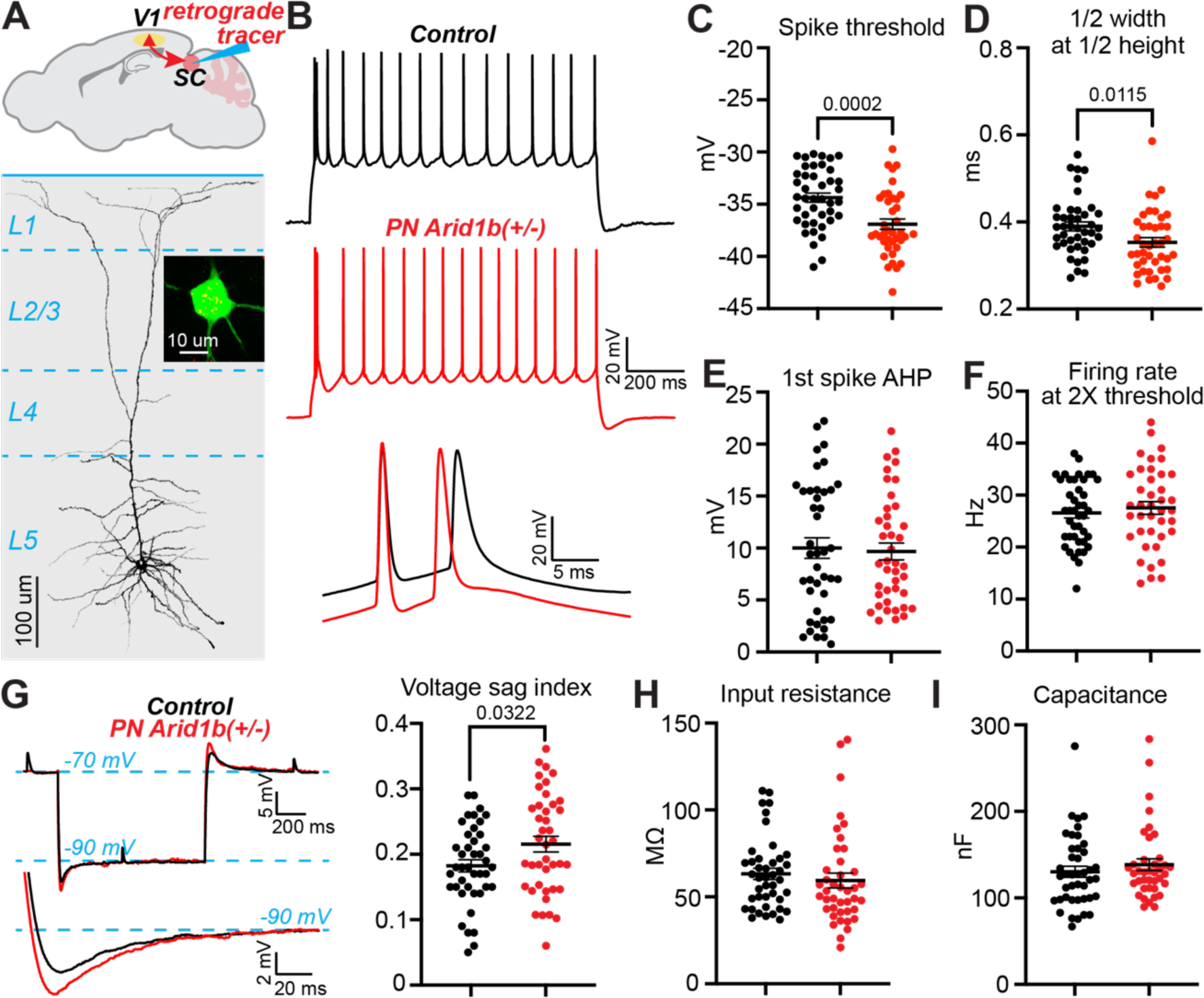
Arid1b haploinsufficiency in layer 5 corticofugal pyramidal cells causes changes in membrane properties like those observed in layer 2/3 fast adapting cells. **A)** (top) Schematic of retrograde tracer injection site in the superior colliculus (SC) to label layer 5 pyramidal-tract type PNs in V1. (bottom) Reconstruction of dendrites traced from biocytin-filled cell. Inset shows biocytin (green) with red retrobeads (yellow) for traced cell. **B)** Examples of responses of pyramidal-tract type PNs in control and PN *Arid1b*(+/-) mice in response to a constant amplitude current pulse. Note hyperpolarized action potential threshold in bottom expansion. **C)** Action potential voltage threshold. Control n = 42, PN Arid1b(+/-) n = 39; unpaired t test. **D)** Half-width at half-height for the first action potential. Control n = 42, PN Arid1b(+/-) n = 41; Mann-Whitney U test. **E)** Afterhyperpolarization amplitude after the first action potential. Control n = 42, PN Arid1b(+/-) n = 41; p = 0.9945, Mann-Whitney test. **F)** Firing rate at twice action potential threshold. Control n = 42, PN Arid1b(+/-) n = 40; p = 0.5317, t test. **G)** Hyperpolarization-induced membrane potential voltage sag. Control n = 42, PN Arid1b(+/-) n = 41; unpaired t test. **H)** Input resistance. Control n = 42, PN Arid1b(+/-) n = 41; p = 0.1541, Mann-Whitney U test. **I)** Membrane capacitance. Control n = 42, PN Arid1b(+/-) n = 39; p = 0.3624, Mann-Whitney U test.

### *Arid1b* haploinsufficiency in pyramidal neurons results in increased local recurrent excitation in layer 2/3

Previous studies recorded spontaneous miniature excitatory postsynaptic currents (mEPSCs) from PNs in mice with whole-body *Arid1b* haploinsufficiency and reported conflicting results; Kim et al. (2022) found a reduction in mEPSC frequency while Jung et al. (2017) did not. Thus, we next investigated if *Arid1b* haploinsufficiency affects synaptic connectivity and physiology among PNs in layer 2/3 by performing paired whole-cell patch clamp recordings between these cells in control and PN *Arid1b*(+/-) mice (**Fig. 4A**). During recording, we observed a mix of CA and FA PN types that formed synaptic connections both within and across types, thus we pooled the data. In control mice, we observed a low connectivity rate between neighboring PNs (**Fig. 4B**), consistent with previous studies (Rinaldi et al., 2008; Brown and Hestrin, 2009; Patel et al., 2014). Surprisingly, in PN *Arid1b*(+/-) mice the connectivity rate was higher (**Fig. 4B**), suggesting abnormal local hyperconnectivity that may increase recurrent excitation within layer 2/3. Although these data seem at odds with the previous studies (Jung et al., 2017; Kim et al., 2022), they are consistent with observations from multiple mouse models of autism that neighboring PNs may be hyperconnected, but excitatory afferent input from other sources is reduced (Courchesne and Pierce, 2005; Rinaldi et al., 2008; Markram and Markram, 2010; Patel et al., 2014; Antoine et al., 2019). Our data suggest a similar circuit phenotype in the *Arid1b* haploinsufficiency model.

**Figure 4.**
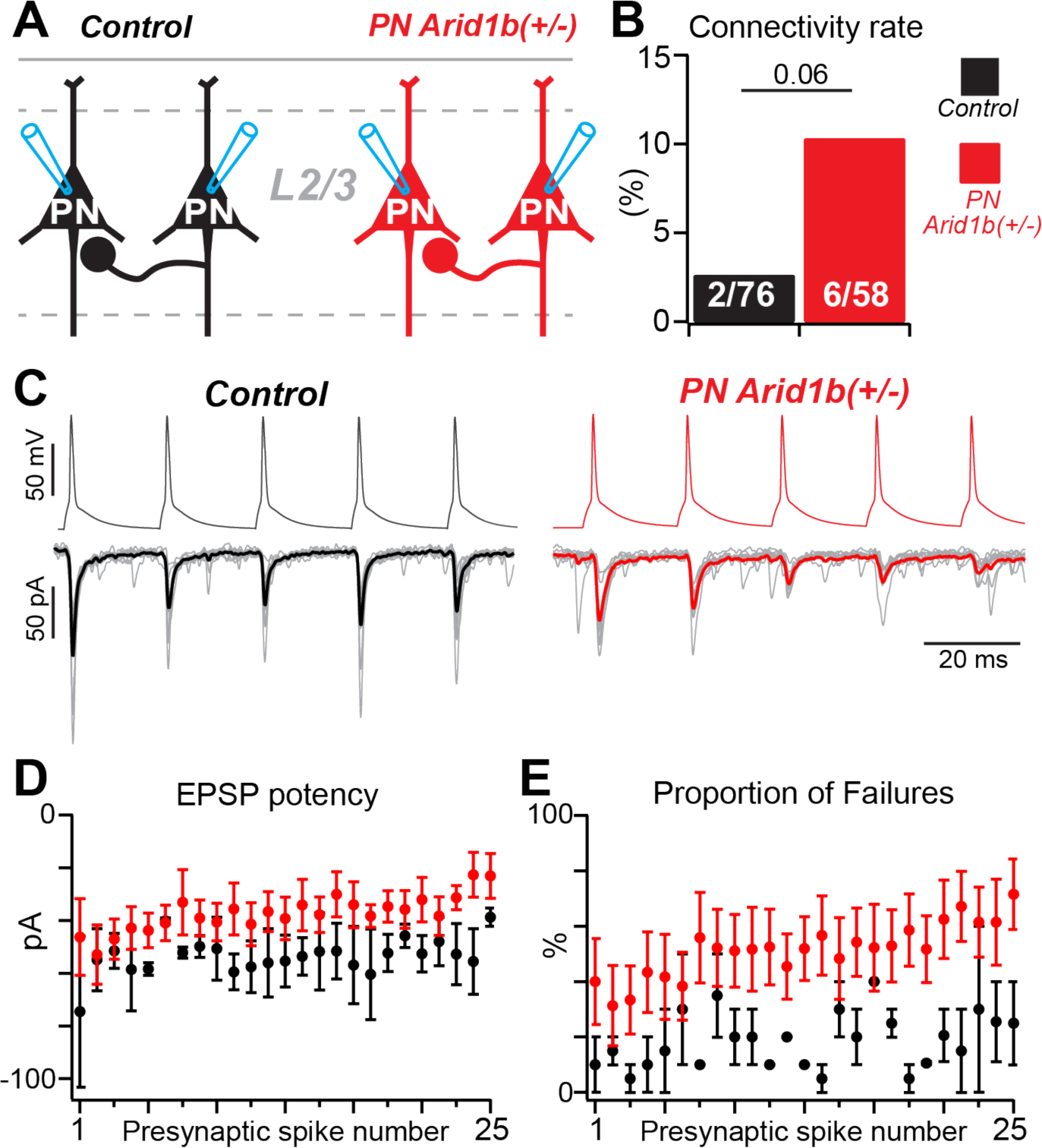
Arid1b haploinsufficiency results in increased synaptic connectivity rate between pyramidal neurons in layer 2/3. **A)** Schematic of paired whole-cell recording configuration for connectivity and physiology of synapses between PNs in layer 2/3. **B)** Synaptic connectivity rates for paired recordings between PNs. Chi-squared test (df = 1). **C)** Examples of unitary excitatory postsynaptic currents in paired recordings. Only the first 5 spikes of the 25-spike train shown. Gray traces are synaptic currents from individual trials; bold black and red lines are average synaptic currents. **D)** Potency of excitatory synapses between PNs. **E)** Percentage of trials in which a presynaptic action potential failed to evoke vesicle release for a connected pair of PNs.

To analyze synaptic physiology, we evoked trains of 25 action potentials in the presynaptic neuron while recording the postsynaptic neuron in voltage clamp with a holding potential of -70 mV for 10 trials (**Fig. 4C**). Synapses between PNs were qualitatively similar between control and PN *Arid1b*(+/-) mice, demonstrating synaptic depression during trains of presynaptic action potentials (**Fig. 4C**). To compare synaptic properties between conditions, we analyzed synaptic potency (average EPSC amplitude for trials in which a presynaptic action potential evoked vesicle release) and synaptic failure rate (percentage of trials in which a presynaptic action potential did not evoke vesicle release) (Stevens and Wang, 1995). Because synaptic connections were rare (n = 2 for control, n = 6 for PN *Arid1b*(+/-) mice) we did not run statistical tests between conditions. Qualitatively, synaptic potency was similar between conditions, however, we noted it was smaller PN *Arid1b*(+/-) mice (**Fig. 4D**). Furthermore, the synaptic failure rate was higher in PN *Arid1b*(+/-) mice (**Fig. 4E**). Although it is difficult to draw conclusions from these limited data points, the observation that excitatory synapses are weaker in PN *Arid1b*(+/-) mice is consistent with our findings below for which we have greater statistical power. Furthermore, they are consistent with previous work reporting reduced synapse density in PNs (Ka et al., 2016; Kim et al., 2022).

### Enhanced strength of inhibitory synapses from PV+ interneurons to pyramidal neurons may rebalance the ratio of excitation-to-inhibition in layer 2/3

Our data suggest at least two potential sources of increased excitation-to-inhibition in PN *Arid1b*(+/-) mice: hyperpolarized spike threshold and increased recurrent excitation among PNs. However, it is unknown if *Arid1b* haploinsufficiency in PNs impacts their synaptic connectivity with neighboring inhibitory interneurons (INs) or the physiology of these synapses, which could either further imbalance the ratio of excitation-to-inhibition or counterbalance it. We chose to investigate synapses to and from parvalbumin (PV) expressing INs, which provide powerful perisomatic inhibition of PNs (Hu et al., 2014). To target PV INs for paired whole-cell recordings with PNs in layer 2/3, we crossed control and PN *Arid1b*(+/-) mice to a PV-tdTomato reporter line (Kaiser et al., 2016) (**Fig. 5A**). In these experiments, we used an internal solution with a chloride reversal potential of -29 mV when patching PNs and -79 mV when patching PV INs. This allowed us to render both glutamatergic and GABAergic currents inward at a holding potential of -70 mV to investigate reciprocal unitary synaptic connections between pairs of neurons (**Fig. 5A, bottom**).

**Figure 5.**
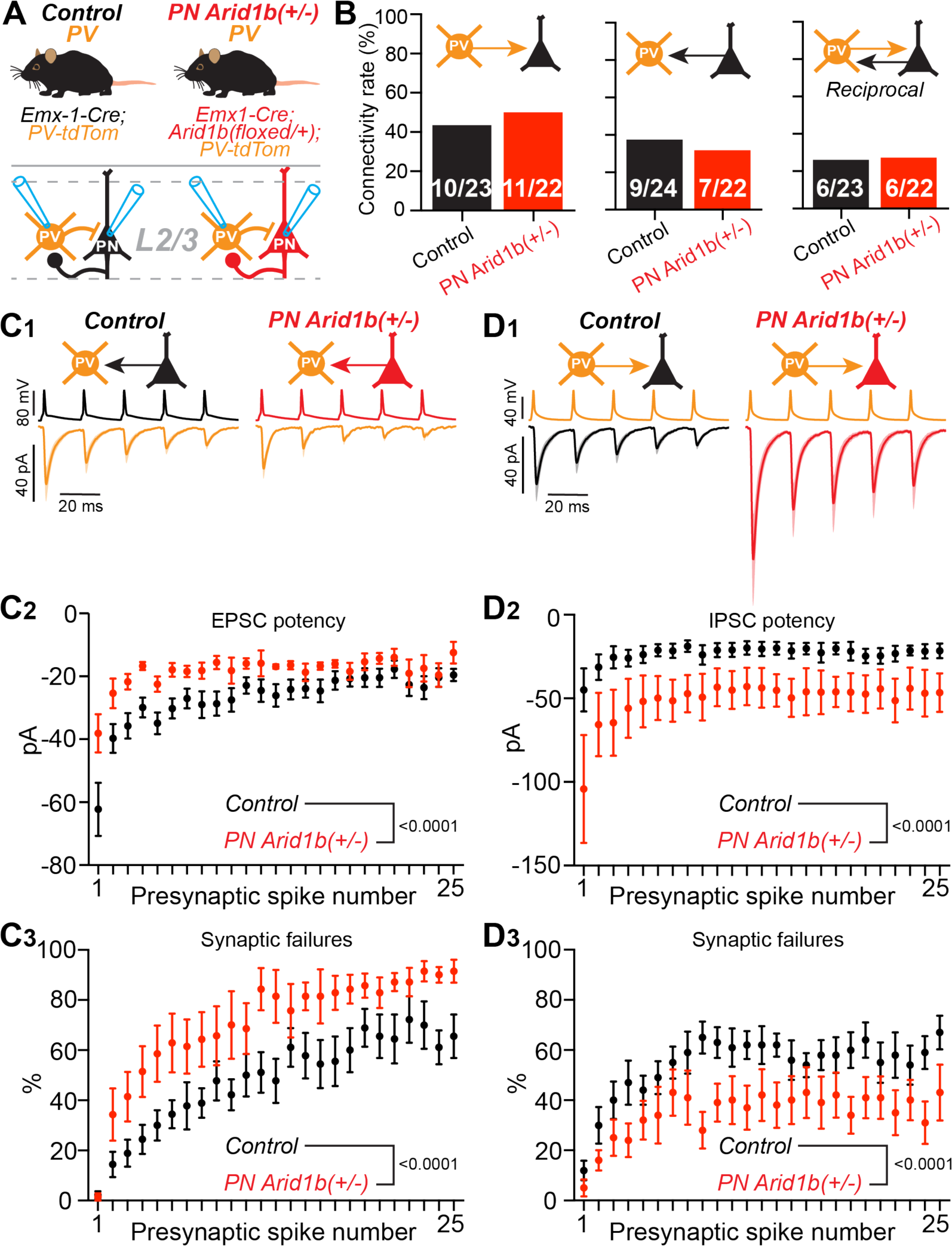
*Arid1b* haploinsufficiency in pyramidal neurons weakens excitatory synaptic input to PV+ interneurons, but PV+ interneurons compensate with stronger inhibitory synapses. **A)** Schematic of paired whole-cell recording configuration for connectivity and physiology of synapses between PNs and PV INs in layer 2/3 and mouse genotypes for each condition. **B)** Synaptic connectivity rates for paired recordings between PNs and PV INs. (left) PV INs to PNs, (middle) PNs to PV INs, (right) reciprocal connections. **C)** (1) Average excitatory postsynaptic currents recorded in PV INs (+/- SEM). Control n = 9, PN Arid1b(+/-) n = 7 connected pairs; (2) Potency of excitatory synapses from PNs to PV INs. Main effect of genotype, two-way ANOVA: F(1,300) = 63.79. (3) Percentage of trials in which a presynaptic action potential in a PN failed to evoke vesicle release at synapse onto a PV IN. Main effect of genotype, two-way ANOVA: F(1,350) = 94.59. **D)** (1) Average inhibitory postsynaptic currents recorded in PNs (+/- SEM). Control n = 10, PN Arid1b(+/-) n = 11 connected pairs; (2) Potency of inhibitory synapses from PV INs to PNs. Main effect of genotype, two-way ANOVA: F(1,436) = 77.46. (3) Percentage of trials in which a presynaptic action potential in a PV IN failed to evoke vesicle release at synapse onto a PN. Main effect of genotype, two-way ANOVA: F(1,450) = 74.70.

We found that *Arid1b* haploinsufficiency in PNs did not alter the probability of observing synaptic connections from PV INs to PNs (**Fig. 5B, left**) or PNs to PV INs (**Fig. 5B, middle**), or the probability of observing reciprocally connected pairs (**Fig. 5B, right**). However, we observed striking differences in the physiology of excitatory synapses from PNs to PV INs and inhibitory synapses from PV INs to PNs. For synaptic connections from PNs to PV INs, the average amplitude of EPSCs was smaller in PN *Arid1b*(+/-) mice (**Fig. 5C1**). To quantify changes in synaptic physiology, we analyzed the potency and failure rate of excitatory synapses onto PV INs, as described above for PN-to-PN synapses (**Fig. 4D, E**). Interestingly, excitatory synaptic potency was lower (**Fig. 5C2**) and synaptic failure rate was higher (**Fig. 5C3**) in PN *Arid1b*(+/-) mice relative to control. Thus, *Arid1b* haploinsufficiency in PNs weakens local excitatory drive onto PV INs, which may exacerbate excitation-to-inhibition imbalance. However, surprisingly, for synaptic connections from PV INs to PNs, the average amplitude of inhibitory postsynaptic currents was larger in PN *Arid1b*(+/-) mice (**Fig. 5D1**). This was due to increased potency of inhibitory currents (**Fig. 5D2**) and reduced synaptic failure (**Fig. 5D3**) rate relative to controls. Importantly, in these experiments PV INs were genetically normal. Thus, it is possible that increased inhibitory strength in PN *Arid1b*(+/-) mice occurs due to do homeostatic plasticity mechanisms to maintain network activity stability (McFarlan et al., 2023). However, we cannot rule out the possibility that changes at this synapse are due directly to *Arid1b* haploinsufficiency in PNs and thus “pathological”. Regardless, enhanced inhibition of PNs in PN *Arid1b*(+/-) mice may compensate for weak excitation of PV INs, increased recurrent excitation among PNs, and hyperpolarized spike threshold in PNs to rebalance the ratio of excitation-to-inhibition.

### *Arid1b* haploinsufficiency in pyramidal neurons is insufficient to cause behavioral deficits or seizure vulnerability observed in the preclinical mouse model

Preclinical mouse models of *Arid1b* haploinsufficiency, in which all cells of the body lack one functional copy of the gene, demonstrate several phenotypes consistent with autism spectrum disorder (ASD), including deficits in social behaviors (Celen et al., 2017; Jung et al., 2017; Shibutani et al., 2017; Ellegood et al., 2021; Kim et al., 2022).

Previously, Jung et al. (2017) found that conditional *Arid1b* haploinsufficiency in inhibitory interneurons was sufficient to generate these behavioral abnormalities. In a follow up study, Moffat et al. (2021b) characterized behavioral phenotypes in PN *Arid1b*(-/-) mice and observed deficits in novel object recognition and social novelty. This suggests cell-autonomous pathology in PNs also contributes to ASD symptoms. However, no studies have investigated if *Arid1b* haploinsufficiency in PNs is sufficient to cause phenotypic changes consistent with ASD.

We compared behaviors between PN *Arid1b*(+/-), whole-body *Arid1b*(+/-), and control mice (**Fig. 6A**). To generate whole-body *Arid1b*(+/-) mice, we crossed Sox2Cre mice (Hayashi et al., 2002; Hayashi et al., 2003) to the same conditional *Arid1b* line used to produce PN Arid1b(+/-) mice. Offspring of female Sox2Cre mice inherit maternal germline Cre recombination during embryonic development, regardless of whether they inherit the Cre allele; we termed these mice germline *Arid1b*(+/-). We also used two sets of control mice for behavioral testing: 1) C57BL/6J mice, on which we ran behavioral tests concurrently with germline *Arid1b*(+/-) mice and 2) *Arid1b*(flox/flox) mice that were not crossed to a Cre transgenic line, on which we ran behavioral tests concurrently with PN *Arid1b*(+/-) mice. This allowed us to confirm that baseline behaviors were stable between germline and PN *Arid1b*(+/-) cohort tests and that Arid1b(flox/flox) mice are behaviorally equivalent to control C57BL/6J mice. First, we tested novel object recognition and found that mice in all conditions spent more time investigating the novel object relative to the familiar with no differences among groups (**Fig. 6B**). Furthermore, in a three-chamber sociability assay in which an empty wire cup was placed on one side and a novel conspecific mouse on the other, mice in all conditions were equally biased towards spending time with a novel conspecific (**Fig. 6C**). However, when given free access to a novel versus familiar conspecific mouse to assess preference for social novelty, only germline *Arid1b*(+/-) mice were biased towards the familiar conspecific; the behavior of PN *Arid1b*(+/-) mice was comparable to both control conditions (**Fig. 6D)**. Thus, our data show that *Arid1b* haploinsufficiency in PNs alone is insufficient to drive deficits in social behavior.

**Figure 6.**
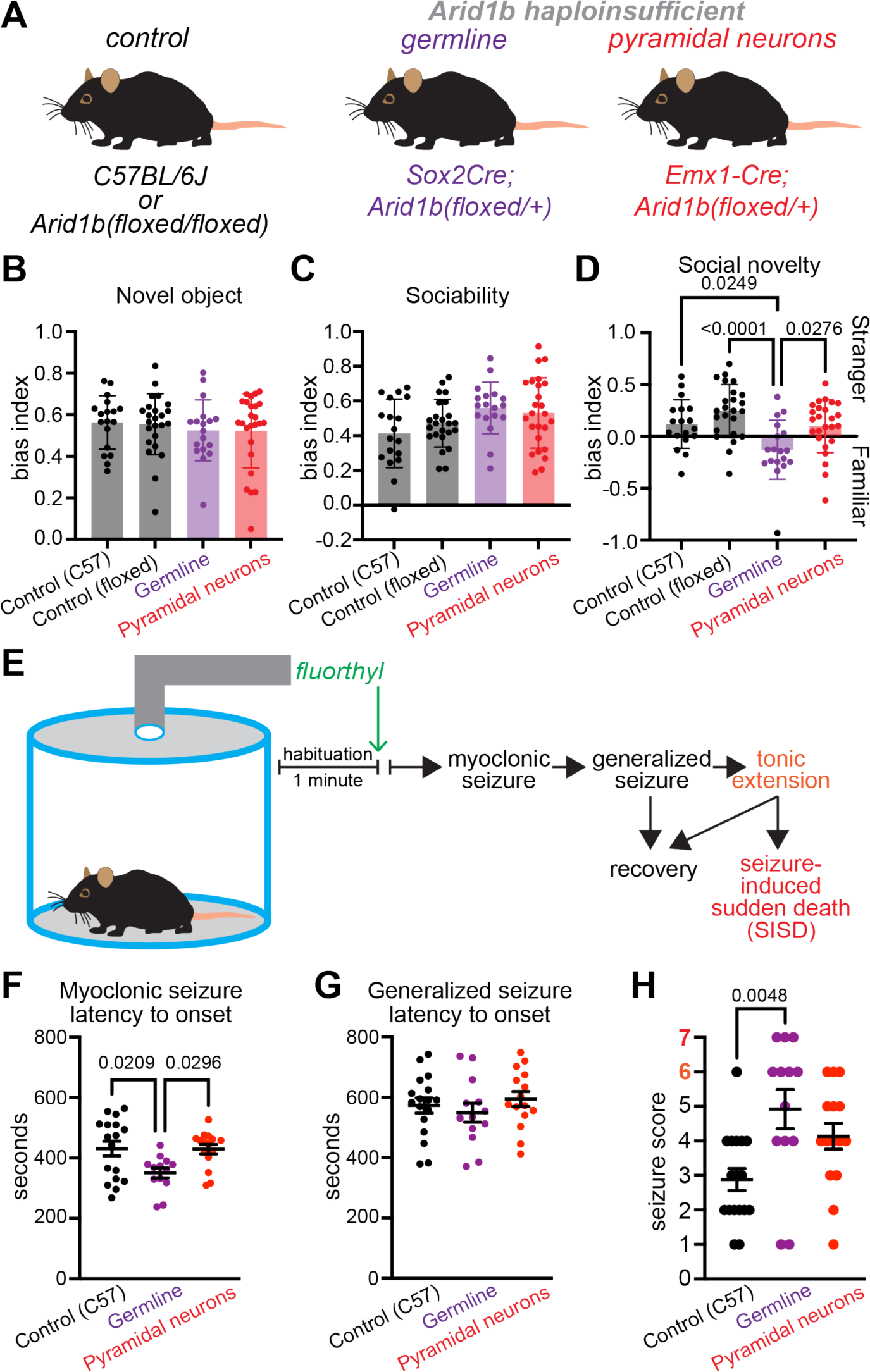
Germline Arid1b haploinsufficiency causes a social novelty deficit and vulnerability to seizures that are not observed in mice with conditional Arid1b haploinsufficiency in pyramidal neurons. **A)** Mouse genotypes for conditions. **B)** Novel object recognition test. Control (C57) n = 18, Control (floxed) n = 24, Germline n = 18, PN Arid1b(+/-) n = 25. Kruskal-Wallis test: H(3) = 1.125, p = 0.7712. **C)** Three-chamber test for sociability. One-way ANOVA: F(3,81) = 2.611, p = 0.0570. n’s same as (B). **D)** Three-chamber test for social novelty. One-way ANOVA: F(3,81) = 7.27, p = 0.0002; Tukey’s post hoc: Control (57) vs Control (floxed) p = 0.411, Control (57) vs. PN Arid1b(+/-) p = 0.9934, Control (floxed) vs. PN Arid1b(+/-) p = 0.2035. n’s same as (B). **E)** Schematic for flurothyl-induced seizure assay and progression of seizure outcomes. 0**F)** Latency to myoclonic seizure onset. One-way ANOVA: F(2,42) = 4.793, p = 0.0133; Tukey’s post hoc: Control (C57) vs. PN Arid1b(+/-) p = 0.9974. Control (C57) n = 17 mice, Germline n = 13 mice, PN Arid1b(+/-) n = 15 mice. **G)** Latency to generalized seizure. One-way ANOVA: F(2,42) = 0.6425, p = 0.5309. n’s same as (F). **H)** Seizure outcome scores. Kruskal-Wallis test: H(2) = 10.52, p = 0.0052; Control (C57) vs. PN Arid1b(+/-) p = 0.1125, PN Arid1b(+/-) vs. PN Arid1b(-/-) p = 0.7805. n’s same as (F).

Finally, human patients with *Arid1b* haploinsufficiency have increased susceptibility to seizures, suggesting hyperexcitability of neural circuits (van der Sluijs et al., 2019; Proietti et al., 2021). Consistent with human studies, previous work found that mice with whole-body *Arid1b* haploinsufficiency exhibit higher susceptibility to seizures induced by a lethal dose of pentylenetetrazol (Ellegood et al., 2021). Here, we used a flurothyl vapor assay (Krasowski, 2000; Kadiyala et al., 2014; Judson et al., 2016) to compare seizure threshold and severity among control (C57BL/6J), germline *Arid1b*(+/-), and PN *Arid1b*(+/-) mice (**Fig. 6E)**. Consistent with Ellegood et al. (2021), we found that germline *Arid1b*(+/-) mice were more vulnerable to seizures, however, pathology in PNs was insufficient to cause this phenotype. Germline *Arid1b*(+/-) mice had a reduced latency to myoclonic seizures relative to both control and PN *Arid1b*(+/-), which were comparable (**Fig. 6F)**. Furthermore, although the latencies to generalized seizure were comparable among conditions (**Fig. 6G)**, germline *Arid1b*(+/-) mice exhibited the most severe seizure outcomes (Samoriski and Applegate, 1997) (**Fig. 6H)**. Over half of germline *Arid1b*(+/-) mice experienced tonic limb extension (7/13 with seizure score >= 6), and three died immediately after (seizure score = 7) (**Fig. 6H)**. In contrast, none of the PN *Arid1b*(+/-) died and most had seizure scores below 6 (12/15 with seizure score <= 5) (**Fig. 6H)**. However, we note seizure scores for PN *Arid1b*(+/-) were the most broadly distributed, rendering them not significantly different from either control or germline *Arid1b*(+/-) mice. It is possible that pathology in PNs with *Arid1b* haploinsufficiency contributes to network hyperexcitability; however, it is not sufficient to produce vulnerability to seizures observed in the preclinical mouse model.

## DISCUSSION

### Excitatory projection neuron survival and differentiation as function of *Arid1b* **gene dosage**

ARID1B is a subunit of the Brg/Brahma-associated factor (BAF) chromatin remodeling complex that contains an AT-rich DNA binding domain (ARID) necessary to regulate transcription (Wang et al., 1996; Wang et al., 2004). Because ARID1B controls expression of many genes starting during early development in several tissues (Celen et al., 2017; Moffat et al., 2021a; Pagliaroli and Trizzino, 2021), understanding its function and how its loss causes disease is challenging. However, it is broadly known to play roles in cell proliferation, differentiation, and survival (Yan et al., 2008; Son and Crabtree, 2014; Pagliaroli et al., 2021; Pagliaroli and Trizzino, 2021). In previous work, Moffat et al. (2021b) used the Emx1-IRES-Cre mouse line to conditionally knock out both copies of *Arid1b* from dorsal progenitors, which includes excitatory projection neurons. They found that complete loss of ARID1B resulted in an increase in the number of progenitors undergoing apoptosis and a concomitant reduction in the number of intermediate progenitor cells. In juvenile mice, they found fewer Cux1+ cells in cortical superficial layers, indicating a subset of excitatory projection neurons were lost. However, it was unclear why some progenitors were uniquely vulnerable to ARID1B loss.

Here, we used the same transgenic strategy as Moffat et al. (2021b) to conditionally knockout either one or both copies of *Arid1b*, and then investigated the impact on excitatory neuron physiology in mature mice. Importantly, we found that changes in neuronal survival and membrane properties depend on both cell-type and gene dosage. In control mice, we were able to identity two different types of excitatory projection neurons in layer 2/3 that we termed continuous adapting (CA) and fast adapting (FA) based on their firing responses to constant amplitude depolarizing current pulses. Strikingly, complete knockout of *Arid1b* resulted in the observation of significantly fewer FA cells relative to CA cells. The intrinsic membrane properties of CA cells were mostly unaffected, and changes did not include those that distinguish them from FA cells, such as firing rate, input resistance, and membrane capacitance. Thus, the reduced number of cells classified as FA in PN *Arid1b*(-/-) mice was not due to emergence of a new “hybrid” excitatory subtype in layer 2/3. Rather, our data suggest that the proliferation and/or survival of FA cells, but not CA cells, is uniquely dependent on ARID1B. Finally, although *Arid1b* haploinsufficiency did not affect the proliferation or survival of FA cells, it did alter their intrinsic membrane properties, including hyperpolarization of spike threshold. In contrast, *Arid1b* haploinsufficiency had no effect on the physiology of CA cells. Thus, FA cells, but not CA cells, require ARID1B for their proper development and are vulnerable to changes in *Arid1b* gene dosage.

It is perhaps surprising that loss of *Arid1b* expression, which leads to reduced cortical progenitor production, impacts the survival of only a specific subset of excitatory projection neurons. However, these data are congruent with results from experiments on the effects of *Arid1b* loss from ventral progenitors in the medial ganglionic eminence (MGE) (Jung et al., 2017; Moffat et al., 2021b). The MGE is a transient progenitor pool that gives rise to inhibitory interneurons subtypes distinguished by expression of parvalbumin (PV) and somatostatin (SST) (Butt et al., 2005; Rudy et al., 2011). Interestingly, a single MGE progenitor can give rise to both PV+ and SST+ subtypes (Brown et al., 2011; Ciceri et al., 2013; Harwell et al., 2015; Mayer et al., 2015). However, loss of *Arid1b* leads to a reduction of PV+ but not SST+ cells in mature mice (Jung et al., 2017; Moffat et al., 2021b). Our data suggest that a similar cell-type-specific vulnerability exists for dorsal progenitors.

### Changes in excitatory projection neuron physiology in *Arid1b* haploinsufficiency and in other mouse models of ASD

Our study is the first to systematically evaluate the effects of *Arid1b* haploinsufficiency on the intrinsic membrane properties of different subtypes of excitatory projection neurons. We found hyperpolarization of the voltage threshold for spike initiation, reduced spike width, and increased voltage sag in both layer 2/3 FA cells and layer 5 pyramidal-tract type projection neurons. Previously, Kim et al. (2022) examined the physiology of layer 2/3 projection neurons in medial prefrontal cortex (mPFC) at a similar development stage in germline *Arid1b* haploinsufficient mice. However, they did not find a change in the voltage threshold for spike initiation. This may reflect regional differences (mPFC versus V1 in our study) in projection neuron vulnerability to *Arid1b* haploinsufficiency. However, it is also possible that, as in V1, only a specific subset of layer 2/3 cells in mPFC are affected, but these were not sampled during recording. Furthermore, they did not characterize the electrophysiological properties of neurons in layer 5. Future studies are necessary to determine if our results generalize across cortical regions.

ARID1B shares molecular targets with CHD8, another chromatin modifier strongly associated with ASD, and FMRP, which causes Fragile X Syndrome (FXS), and (Darnell et al., 2011; Shibutani et al., 2017; Kim et al., 2022). Interestingly, some changes in intrinsic membrane properties we observed in the *Arid1b* haploinsufficiency mouse model are also found in FXS mice. For example, Kalmbach et al. (2015) observed hyperpolarized spike threshold for layer 5 pyramidal tract-type cells, and Routh et al. (2017) observed reduced spike width in layer 2/3 cells in mPFC. However, while we found increased voltage sag in both layer 2/3 FA cells and layer 5 pyramidal-tract type cells, Kalmbach et al. (2015) found reduced voltage sag and H-current in FXS mice. Furthermore, in a *Chd8* mouse model, Ellingford et al. (2021) found no changes in the voltage threshold or width of spikes of layer 5 excitatory cells in mPFC. Thus, our data suggest some changes in neuronal physiology converge in *Arid1b* haploinsufficiency and FXS, but future studies are needed to directly compare the same neuronal subtypes in the same cortical regions.

### Altered excitatory synaptic connectivity and physiology in *Arid1b* **haploinsufficiency is similar to FXS**

We used paired whole-cell recordings to study if *Arid1b* haploinsufficiency in excitatory projection neurons alters the synaptic physiology of unitary connections they send and receive with neighboring projection neurons and PV+ inhibitory interneurons. Interestingly, although we observed increased connectivity rate among layer 2/3 projection neurons, we also found decreases in the strength of excitatory synapses to both excitatory and inhibitory cells. Our data are at odds with previous work that recorded spontaneous miniature excitatory postsynaptic currents (mEPSCs) from excitatory cells in mPFC. Specifically, Kim et al. (2022) found reduced frequency but not amplitude of mEPSCs, while Jung et al. (2017) found no differences. However,mechanisms that regulate evoked versus spontaneous neurotransmitter release are distinct (Ramirez and Kavalali, 2011), and discrepancies between synaptic connectivity rate measured from paired recordings and spontaneous mEPSC frequency have been observed previously in studies of FXS mice (Gibson et al., 2008). Interestingly, our evoked synaptic data are broadly consistent with those observed in the FXS model. Specifically, in layer 5A of somatosensory cortex, which is populated by intratelencephalic-type projection neurons, excitatory synaptic pruning during development is impaired, leading to an increased connectivity rate between cells (Patel et al., 2014). A similar mechanism may occur in *Arid1b* haploinsufficient mice. Furthermore, multiple studies found reduced strength of excitatory synaptic input to PV+ interneurons in FXS mice (Gibson et al., 2008; Patel et al., 2013; Nomura et al., 2017; Antoine et al., 2019). In particular, Patel et al. (2013) found that this is due to pathology in the presynaptic excitatory neurons, which leads to reduced evoked transmitter release. Our cellular and synaptic physiology data in *Arid1b* haploinsufficient mice are surprisingly congruent with potential pathophysiological mechanisms underlying FXS.

### Contribution of cell-autonomous pathology in excitatory neurons to ASD behavioral and seizure phenotypes

We found that inducing conditional *Arid1b* haploinsufficiency in all dorsal cortical progenitors was insufficient to cause deficits in social behavior or susceptibility to seizures observed in a preclinical germline model. These data are consistent with recent work that strongly implicate pathology in cortical inhibitory interneurons as a major driver of ARID1B related disorders (Jung et al., 2017; Smith et al., 2020; Moffat et al., 2021b) and ASD more broadly (Judson et al., 2016; Lunden et al., 2019; Contractor et al., 2021; Nomura, 2021; Tang et al., 2021). However, pathology in neurons outside the cortex may play an important role in *Arid1b* haploinsufficiency phenotypes. For example, conditional deletion of diverse ASD risk genes from peripheral sensory neurons also results in deficits in social behavior, including social novelty (Orefice et al., 2016; Orefice et al., 2019). Furthermore, conditional deletion of *Fmr1* from glutamatergic neurons of the inferior colliculus but not cortical projection neurons results in susceptibility to audiogenic seizures (Gonzalez et al., 2019). In conclusion, our data support a growing body of literature suggesting, rather surprisingly, that pathology in cortical excitatory projection neurons plays a relatively minor role in causing ASD.

## METHODS

### Animals

All experiments were conducted in accordance with animal protocols approved by the Ohio State University IACUC. Male and female mice, 5 – 8 weeks old, were used without bias. Emx1-IRES-Cre, Arid1b(floxed/floxed), Sox2Cre, Parvalbumin-tdTomato, and C57BL/6J mice were obtained from the Jackson Laboratory (stock nos. 005628, 032061, 008454, 027395, 000664).

### Slice preparation

Mice were anesthetized with isoflurane and then decapitated. The brain was dissected in ice-cold artificial cerebrospinal fluid (ACSF) containing (in mM): 100 sucrose, 80 NaCl, 3.5 KCl, 24 NAHCO3, 1.25 NaH2PO4, 4.5 MgCl, 0.5 CaCl2, and 10 glucose, saturated with 95% O_2_ and 5% CO_2_. Coronal slices of primary visual cortex (300 μm) were cut using a Leica VT 1200S vibratome (Leica Microsystems) and incubated in the above solution at 35 °C for 30 minutes post-dissection. Slices were then maintained at room temperature until use in recording ACSF containing (in mM): 130 NaCl, 3.5 KCl, 24 NaHCO3, 1.25 NaH2PO4, 1.5 MgCl, 2.5 CaCl2, and 10 glucose, saturated with 95% O_2_ and 5% CO_2_.

### Electrophysiology

For recording, slices were constantly perfused with ACSF at 2 mL/min at a temperature of 31-33 °C. Cells were visualized using an upright microscope (Scientifica SliceScope) with a 40x water-immersion objective (Olympus), camera (Scientifica SciCam Pro), and Oculus software. Retrobeads and tdTomato-expression were identified by fluorescence (CoolLED pE-300ultra). Recording pipettes were pulled from borosilicate glass (World Precision Instruments) to a resistance of 3-5 MΩ using a vertical pipette puller (Narishige PC-100). Whole-cell patch clamp recordings were acquired using a Multiclamp 700B amplifier (Molecular Devices), filtered at 3 kHz (Bessel filter), and digitized at 10 or 20 kHz (Digidata 1550B and pClamp v11.1, Molecular Devices). Recordings were not corrected for liquid junction potential. Series resistance (10-25 MΩ) was closely monitored, and recordings were excluded from analysis if series resistance passed 25 MΩ. In current-clamp mode, cells were biased to a membrane potential (Vm) of -70 mV; in voltage-clamp mode, a holding potential of -70 mV was applied. The internal solution used to collect intrinsic membrane properties and excitatory postsynaptic currents and potentials contained (in mM): 130 K-gluconate, 5 KCl, 2 NaCl, 4 MgATP, 0.3 NaGTP, 10 phosphocreatine, 10 HEPES, 0.5 EGTA, and 0.2% biocytin (Sigma-Aldrich catalog #B4261). The calculated ECl-for this solution was -79 mV. The internal solution used to record GABA_A_-mediated currents in voltage-clamp at a holding potential of -70 mV contained (in mM): 85 K-gluconate, 45 KCl, 2 NaCl, 4 MgATP, 0.3 NaGTP, 10 phosphocreatine, 10 HEPES, and 0.5 EGTA, for a calculated ECl-of -29 mV. The pH of both internal solutions was adjusted to 7.4 with KOH.

For paired recordings of synaptic connections, presynaptic cells were made to fire trains of action potentials (25 at 50 Hz) using 2 nA steps of 2 ms duration every 10 s for 10 trials. Post-synaptic cells were recorded in voltage-clamp at -70 mV. To analyze postsynaptic currents in paired recordings, we first zeroed the data by subtracting the baseline and then performed 10 repetitions of binomial (Gaussian) smoothing. All postsynaptic currents (PSCs) were then analyzed relative to the timing of the peak of each presynaptic spike during the train. PSCs were detected by threshold crossing (-7 to -10 pA); if this threshold was already crossed at the instant of spike peak, the PSC data associated with that spike of that trial were discarded as being contaminated by spontaneous events. The proportion of failures was calculated as the ratio of evoked PSCs to the total number of non-contaminated trials for each presynaptic spike. The PSC potency was calculated as the average peak amplitude of all successfully evoked PSCs for each presynaptic spike (i.e., failures were not included in the average).

Input resistance (Rin) was measured using a linear regression of voltage deflections (±20 mV from -70 mV resting membrane potential) in response to 1 s current steps. To calculate voltage sag, the membrane potential was biased to -70 mV (V_initial) followed by injection of a 1 s negative current step of sufficient amplitude to reach a steady state Vm of -90 mV during the last 200 ms of the current injection (V_sag). The peak hyperpolarized Vm prior to sag (V_hyp) was used to calculate the sag index as (V_hyp – V_sag)/(V_hyp – V_initial). Capacitance was calculated as 1/Rin, both measured from the voltage response to a 10 pA current step relative to a -70 mV resting potential. The voltage threshold for action potential initiation was measured from the response to the minimum amplitude current step necessary to generate a spike. Spike threshold was defined as the voltage recorded at the maximum of the second derivative of voltage change with respect to time (Wilent and Contreras, 2005). Spike half-width at half-height was calculated between spike threshold and spike peak.

### Reconstruction of dendritic morphology

Following whole-cell recording with biocytin, tissue slices were drop-fixed for 24 hours in 4% paraformaldehyde. Slices were then rinsed with PBS and incubated in the secondary antibody streptavidin Alexa Fluor 488 (1:1000, Invitrogen of Thermo Fisher Scientific catalog #S32354, RRID: AB_2315383) and DAPI (1:2000, Thermo Fisher Scientific catalog #D1306) overnight at 4 °C. All antibodies were diluted in carrier solution consisting of PBS with 1% BSA, 1% normal goat serum, and 0.5% Triton X-100. Slices were next rinsed with PBS and resectioned on a freezing microtome at 100 μm and slide mounted. Sections were imaged using a Leica DM6 CS with a 20X objective. Reconstructions of dendrites were made using Simple Neurite Tracer (Longair et al., 2011) for Fiji (Schindelin et al., 2012).

### Stereotaxic injections

For injections of retrograde tracer into the superior colliculus, mice were anesthetized with 5% isoflurane and mounted in a stereotax (Neurostar, Germany). 24 hours prior to surgery, ibuprofen was added to the cage drinking water and maintained for 72 hours post-operation. Prior to surgery, mice were provided with extended release (48 – 72 hour) buprenorphine (1 mg/kg via subcutaneous injection) for additional post-operative analgesia. Red retrobeads (Lumafluor) or retrograde serotype AAV-CAG-tdTomato (gift from Edward Boyden; Addgene Addgene viral prep # 59462-AAVrg) were injected via a glass capillary nanoinjector (Neurostar, Germany) back filled with light mineral oil. The superior colliculus was targeted using the following coordinates: 4.0 (±0.2) mm caudal and 0.4 mm lateral to bregma, and 1.6 mm deep from the dura. 100 nl of retrobeads or 400 nl of AAV were injected at 75 nl/min. The pipette was left in place for 10 min following the injection before removal. For retrobead injections, mice were sacrificed after at least 2 days post-injection; for AAV injections, mice were sacrificed after at least 7 days post-injection.

### Behavioral phenotyping

Tests for novel object recognition were performed in a cage with dimensions (in inches) of 10.5 (W) X 14.5 (L) X 9 (H) over four sessions, each lasting 10 minutes. In the first session, mice were allowed to habituate to the testing cage. In the second session, two identical objects were placed in the cage for mice to freely explore. In the third session, one these objects were moved to a new location. Finally, in the fourth session, one of the objects was replaced with a novel object. The novel object bias index was calculated as: ((time exploring novel object) – (time exploring familiar object)) / (total time exploring both) during the fourth session.

Tests for sociability and social novelty were performed in an apparatus with dimensions (in inches) of 7 (W) X 16 (L) X 8 (H) and consisting of three chambers separated by plexiglass dividers. The two end chambers each contained a wire mesh cup. These behavioral assays were conducted over three sessions each lasting 10 minutes. In the first session (habituation), an experimental mouse was placed in the middle chamber, the plexiglass dividers were removed, and the mouse was free to explore the apparatus. The experimental mouse was then returned to the middle chamber and the plexiglass dividers were re-installed. In the second session, to test sociability, an age- and sex-matched C57BL/6J mouse (i.e., conspecific) was placed in one of wire mesh cups, and the plexiglass dividers were removed to allow the mouse to freely explore. The sociability index was calculated as: ((time exploring mouse) – (time exploring empty cup)) / (total time exploring both). The experimental mouse was returned to the middle chamber and the plexiglass dividers were re-installed. In the third session, to test social novelty, a new age- and sex-matched C57BL/6J mouse was placed in the empty wire mesh cup, and the plexiglass dividers were removed to allow the experimental mouse to freely explore a novel or familiar conspecific. The social novelty index was calculated as: ((time exploring novel mouse) – (time exploring familiar mouse)) / (total time exploring both).

### Flurothyl-induced seizure phenotyping

Each mouse was placed in a 2 L glass chamber and allowed to habituate for 1 min before the top of the chamber was closed. 10% flurothyl (bis-2,2,2-trifluoroethyl ether; Sigma-Aldrich) in 95% ethanol was then infused at a rate of 200 µL/min onto a stack of three filter papers suspended at the top of the chamber. In this assay, mice first exhibit a myoclonic seizure, which is defined by transient sudden involuntary jerk/shock-like movements involving the face, trunk, and/or limbs. The intermittent myoclonic seizure progresses to a more sever and continuous generalized seizure. The severity of generalized seizure was scored based on criteria previously described (Samoriski and Applegate, 1997): grade 1, purely clonic seizure; grade 2, “transitional” behaviors involving high-frequency/low-magnitude bouncing and/or rapid backward motion; grade 3, running/bouncing episode; grade 4, secondary loss of posture with bilateral forelimb and hindlimb treading; grade 5, secondary loss of posture with bilateral forelimb tonic extension and bilateral hindlimb flexion followed by treading; grade 6, secondary loss of posture with bilateral forelimb and hindlimb tonic extension followed by treading and recovery; grade 7, tonic extension followed by sudden cardiorespiratory failure and death. Assigned score reflects the highest-grade seizure behavior expressed by the animal within a trial. Upon emergence of a generalized seizure, the lid of the chamber was immediately removed, allowing for rapid dissipation of the flurothyl vapors and exposure of the mouse to fresh air. The flurothyl chamber was recharged with fresh filter paper, cleaned using water, and thoroughly dried between subjects.

### Data analysis and statistics

Electrophysiology data were analyzed in Igor Pro v8.04 (WaveMetrics) using custom routines; pClamp files were imported into Igor using NeuroMatic (Rothman and Silver, 2018). Statistical tests were performed using Graphpad Prism v10.1.1 (Graphpad Software, San Diego, CA). Distributions were first tested for normality using a Shapiro-Wilk test. Distributions that did not violate normality were compared using parametric tests (unpaired t test, one-way ANOVA, or two-way ANOVA as appropriate, described in figure legends), while those that did were compared using nonparametric tests (Mann-Whitney U test or Kruskal-Wallis test as appropriate, described in figure legends). Tests of layer 2/3 PN subtype distributions were performed with a chi-squared test. Post hoc analyses were performed as appropriate and described in figure legends. Data in the text and graphs are reported as mean ± SEM.

## Author contributions

J.C.W. designed the experiments. A.H.M, M.A.H, D.J.B., D.N., N.B., J.F., E.G., B.G., O.N.K-C., and J.C.W. collected and analyzed the data. A.H.M. and J.C.W. wrote the paper.

## Acknowledgements

This work was supported by funding from the Simons Foundation Autism Research Initiative (SFARI Pilot Award #724187) and NIH (R01MH124870) to J.C.W.

